# Premeiotic and meiotic failures lead to hybrid male sterility in the *Anopheles gambiae* complex

**DOI:** 10.1101/507889

**Authors:** Jiangtao Liang, Igor V. Sharakhov

## Abstract

Hybrid male sterility contributes to speciation by restricting gene flow between related taxa. Detailed cytological characterizations of reproductive organs in hybrid males is important for identifying phenotypes that can help guide searches of speciation genes. To investigate possible cellular causes of hybrid male sterility, we performed crosses between closely related species of the *Anopheles gambiae* complex: *An. merus* with *An. gambiae* or *An. coluzzii*. We demonstrate that hybrid male sterility in African malaria mosquitoes involves two defects in the reciprocal crosses: a premeiotic arrest of germline stem cells in degenerate testes and a failure of the reductional meiotic division of primary spermatocytes in normal-like testes. The premeiotic arrest in degenerate testes of hybrids is accompanied by a strong suppression of meiotic and postmeiotic genes. Unlike pure species, sex chromosomes in normal-like testes of F1 hybrids are largely unpaired during meiotic prophase I and all chromosomes show various degrees of insufficient condensation. Instead of entering reductional division in meiosis I, primary spermatocytes prematurely undergo an equational mitotic division producing nonmotile diploid sperm. Thus, our study identified cytogenetic errors in interspecies hybrids that arise during the early stages of postzygotic isolation.

## 1. Introduction

When species diverge, hybrid offspring that are produced can suffer from reduced fitness. Hybrid male fertility is usually one of the first of these postzygotic phenotypes affected [1, 2]. Therefore, the genetic factors, cellular basis, and molecular mechanisms of hybrid male sterility (HMS) are of considerable interest, as they inform our understanding of both speciation and normal fertility function. Studies in animals commonly assess testis shape, size, or weight, as well as sperm morphology, density, or motility to define male sterility phenotypes. However, the cellular basis of HMS is rarely investigated. Available cytological studies of spermatogenesis in sterile hybrids indicate that multiple mechanisms of functional sterility are possible. Sterile hybrids between *Mus musculus domesticus* and *M. m. musculus* have spermatogenic arrest in early meiosis I with disrupted homoeologous chromosome pairing and meiotic sex chromosome inactivation [3, 4]. Studies of failed spermatogenesis in different *Drosophila* hybrids observed arrests at the premeiotic stage [5], reduced chromosome pairing, unequal chromosome segregation in meiosis [6, 7], characteristic spermiogenic arrests [5], spermatid abnormalities [8], and problems in sperm bundling and motility [9, 10]. Detailed analyses of cellular phenotypes in testes of various hybrid organisms could help to establish the order of origin of postzygotic isolating barriers and to guide the identification of speciation genes.

The *Anopheles gambiae* complex consists of at least nine morphologically nearly indistinguishable sibling species of African malaria mosquitoes [11-13]. Genome-based estimations of the age of the *An. gambiae* complex vary from 1.85 [14] to as young as 0.526 million years [15]. Genomic introgression is prevalent in autosomal regions of several species indicating naturally occurring interspecies hybridization [14, 15]. Experimental crosses of species from the *An. gambiae* complex often produce sterile F1 hybrid males, conforming to Haldane’s rule of sterility or inviability of the heterogametic sex [16-20]. Early crossing experiments between members of the complex have found that HMS is associated with various degrees of testes atrophy and underdevelopment of sperm [18]. A large effect of the X chromosome on HMS has been demonstrated for crosses between *An. gambiae* and *An. arabiensis* [16, 21]. However, an introgression of the Y chromosome from *An. gambiae* into the background of *An. arabiensis* has shown no apparent influence on male fertility, fitness, or gene expression [17]. Because of the recent evolution and ease of hybridization, sibling species of the *An. gambiae* complex offer great opportunities to provide insights into the cellular basis and molecular mechanisms of HMS.

Here, we investigate possible cellular causes of male sterility in hybrids between sibling species of the *An. gambiae* complex. We asked three specific questions: (i) What cellular processes are involved in causing infertility in hybrid mosquito males? (ii) Are the defects leading to HMS premeiotic, meiotic, or postmeiotic? and (iii) What cytogenetic errors trigger spermatogenic breakdown? We demonstrate that HMS in malaria mosquitoes involves two cellular defects in reciprocal crosses: premeiotic arrest in germline stem cells and the failure of the reductional meiotic division in primary spermatocytes. Our data suggest that meiotic abnormality in hybrid males stems from the unpairing of the sex chromosomes and insufficient chromatin condensation. Thus, our study identifies cytogenetic errors in hybrids that arise during the early stages of postzygotic isolation.

## 2. Material and Methods

### (a) Mosquito strains and crossing experiments

Laboratory colonies of *An. gambiae* ZANU (MRA-594), *An. coluzzii* MOPTI (MRA-763), *An. coluzzii* MALI (MRA-860), and *An. merus* MAF (MRA-1156) were obtained from the Biodefense and Emerging Infections Research Resources Repository (BEI). To perform interspecies crosses, male and female pupae were separated to guarantee virginity of adult mosquitoes. After the emergence of adults, crossing experiments were performed by combining 30 females and 15 males in one cage. At least two blood meals were fed to females and at least three repeats of each cross were conducted. Male gonads were photographed using an Olympus phase contrast microscope BX41 and UC90 digital camera (Olympus, Tokyo, Japan). We observed the testes of 30 males and took pictures of the testes of five males from each cross. A movie of sperm motility for five hybrid males from each cross and five males of pure species was recorded using a UC90 digital camera (Olympus, Tokyo, Japan).

### (b) Chromosome preparation and fluorescence *in situ* hybridization (FISH)

Preparations of chromosomes, DNA probe labeling, and FISH were performed as previously described [22, 23]. Five DNA probes were used for FISH in this study (supplementary material, table S1). Cytogenetic analyses were performed on at least 10 male individuals of each pure species and of each hybrid. Lengths of well-spread metaphase I chromosomes of each species and of each hybrid were measured using the ruler tool in Adobe Photoshop CS6 (Adobe Inc., San Jose, CA, USA) (supplementary material, table S2). For whole-mount FISH and analysis of chromosome pairing, testes from one-day-old adults of pure species and hybrids were dissected in 1×PBS solution and fixed in 3.7% paraformaldehyde in 1×PBS with 0.1% tween-20 (PBST) for 10 min at room temperature. After adding labeled DNA probes, testes were incubated at 75°C for 5 min (denaturation) and then at 37 °C overnight (hybridization). Testes from 6 individuals of pure species and of hybrids were scanned with a Zeiss LSM 880 confocal laser scanning microscope (Carl Zeiss AG, Oberkochen, Germany) to analyze pairing and unpairing of the sex chromosomes. A total of 418 and 489 nuclei at the early stages of meiotic prophase I were analyzed in pure species and hybrids, respectively (supplementary material, table S3). The percentage of cells with pairing and no pairing of sex chromosomes was used to compare parental species and hybrids. Since the variances for both groups were equal (*s*_*max*_*/s*_*min*_ < 2), a statistical two-sample pooled *t-*test was performed using the JMP 13 software (SAS Institute Inc., Cary, NC, USA). For more details, see the supplementary material and methods.

### (c) RNA extraction and reverse transcription polymerase chain reaction (RT-PCR)

We dissected reproductive organs, which include both testes and male accessary glands (MAGs), from 30 males, only testes or only MAGs from 20 males, and ovaries from 30 females. These mosquitoes were 0-12-hour-old virgin adults of *An. coluzzii* MOPTI, *An. merus* MAF, and interspecies hybrids from crosses ♀*An. coluzzii* MOPTI × ♂*An. merus* and ♀*An. merus* × ♂*An. coluzzii* MOPTI. Total RNA was extracted using a Direct-Zol^™^ RNA MiniPrep Kit (Zymo Research, Irvine, California, US). Two-step RT-PCR was performed to analyze gene expression. cDNA for selected genes (supplementary material, table S1) was generated in a 20-μl reaction containing 2 μl of 10× RT buffer, 2 μl of 0.1-M dithiothreitol (DTT), 1 μl of 50 μM oligo(dT)_20_ primer, 1 μl of 10 mM dNTP mix solution, 1 μl of 25mM MgCl_2_, 1 μl of RNaseOUT^™^(40 U/μl), 1 μl of SuperScript^™^ III RT (200 U/μl), 2 μl of RNA, and 6 μl of DEPC-treated water using the SuperScript^™^ III First-Strand Synthesis System for RT-PCR (Invitrogen, Carlsbad, CA, US). Amplification products were visualized in a 2% agarose gel and photographed under the same parameters for each gene.

## 3. Results

### (a) HMS phenotypes at the cellular level

To obtain interspecies hybrids, we performed reciprocal crosses between *An. merus* MAF and *An. gambiae* ZANU or *An. coluzzii* MOPTI and MALI. Backcrossing of F1 males to parental females resulted in induction of the laying of eggs that did not hatch, which confirmed mating of sterile F1 hybrid males since seminal fluids but not sperm are required to induce oviposition [24]. The observed sterility of hybrid males agrees with Haldane’s rule [20] for the majority of interspecies crosses in the *An. gambiae* complex except for crosses between *An. gambiae* and *An. coluzzii*, which produce fertile hybrids of both sexes [16-19, 25, 26]. To investigate the developmental phenotypes involved in hybrid sterility, we dissected testes from adult males obtained from interspecies crosses and from pure species. Normal testes of pure species have a spindle-like shape (supplementary material, figure S1A). We found obvious differences between testes morphology/size in one interspecies cross versus its reciprocal. Hybrid males from crosses between female *An. merus* and male *An. gambiae* or *An. coluzzii* display normal-like testes (supplementary material, figure S1B). In contrast, F1 males from crosses between female *An. gambiae* or *An. coluzzii* and male *An. merus* show severely underdeveloped (degenerate) testes (supplementary material, figure S1C). We then tested whether normal-like and degenerate testes of interspecies hybrids produce any sperm. Sperm reach maturity within two days after emergence of *Anopheles* adult males. In squashed testes of 2-5-day-old adults of pure species, large amounts of mature spermatozoa with long tails can be seen. After we crushed the testes, spermatozoa with vibrant motility escaped from the ruptures (supplementary material, figure S1A, movie S1). However, mature sperm or sperm motility could hardly be seen in squashed or crushed normal-like testes of 2-5-day-old adult hybrids from crosses when *An. merus* was the mother. Instead, we see fewer spermatids and mostly nonmotile spermatozoa with large heads and often two short tails growing from opposite ends of the head (supplementary material, figure S1B, movie S2). Staining with DAPI identified extended premeiotic and spermatogenic stages that cause delay of spermatogonia amplification, spermatocyte divisions, and spermatid differentiation in the normal-like testes (supplementary material, figure S2). Only undifferentiated round cells could be seen in degenerate testes of hybrids from reciprocal crosses when *An. merus* was the father (supplementary material, figure S1C). Given the small size of the degenerate testes, these round cells may represent germline stem cells. Thus, neither normal-like nor underdeveloped testes of the interspecies hybrids produce mature motile spermatozoa.

### (b) The progress of meiosis in males of pure species

Here, we provide the first description of chromosome behavior during meiosis of pure *Anopheles* species. The chromosome complement of *Anopheles* males consists of three chromosome pairs: two autosomes (2 and 3) and X and Y sex chromosomes. With the help of sex-chromosome-specific fluorescent probes (supplementary material, table S1), we followed the progress of meiosis in *An. gambiae, An. coluzzii*, and *An. merus.* Figure 1 shows normal activities of meiotic chromosomes in the testes of *An. gambiae* ZANU. In primary spermatocytes, all homologous autosomes and sex chromosomes pair and display chiasmata in diplotene/diakinesis of prophase I, they align with each other at the cell equator in metaphase I, and then move from each other in anaphase I. In secondary spermatocytes, sister chromatids of each chromosome align with each other at the cell equator in metaphase II and go to opposite poles of the cell during anaphase II. Meiotic divisions produce spermatids that contain a haploid set of autosomes and either a Y or X chromosome. *An. coluzzii* and *An. merus* males show similar behaviors of chromosomes during meiosis (supplementary material, figures S3, S4).

**Figure 1.**
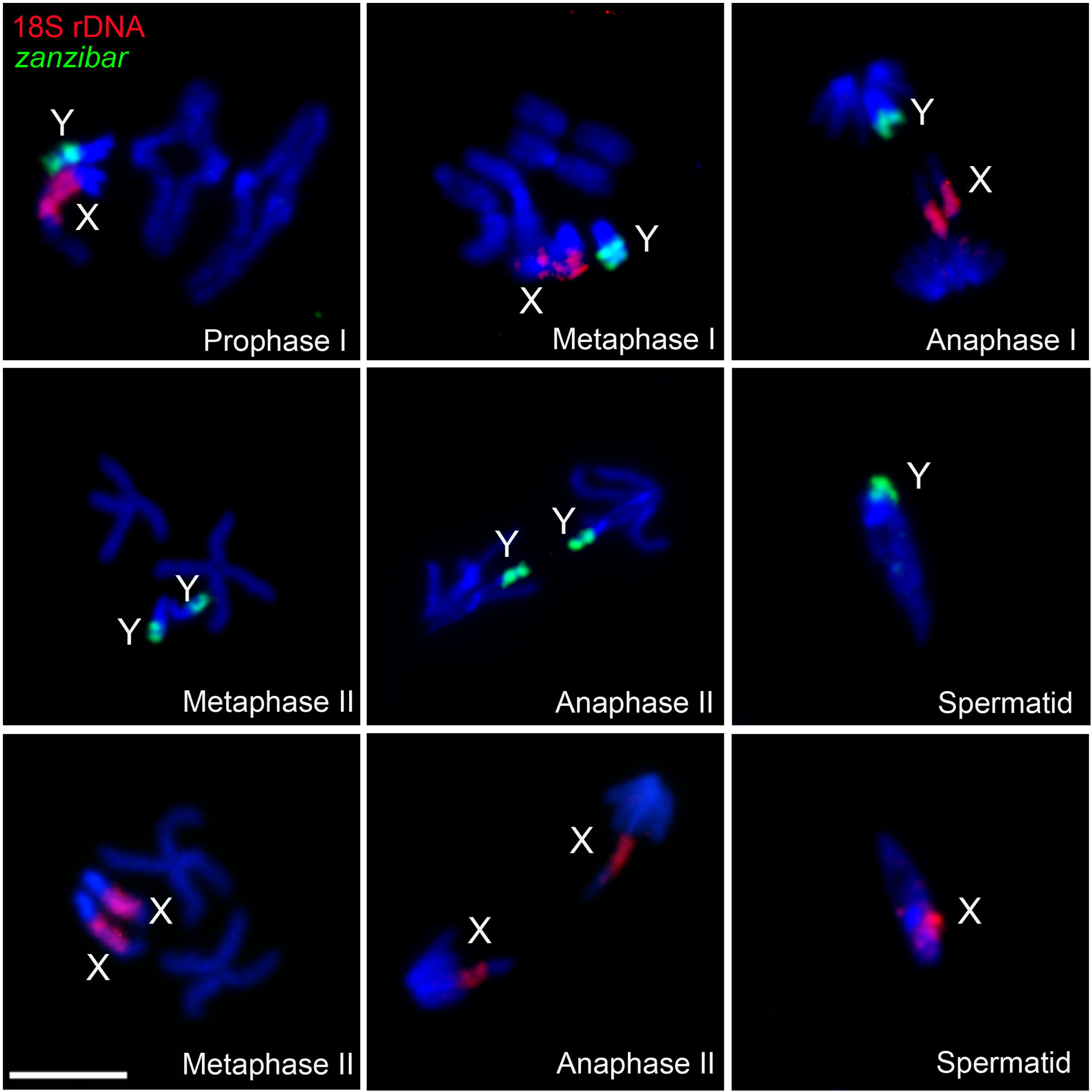
Chromosome behavior during meiosis in testes of *An. gambiae.* X chromosomes are labeled with 18S rDNA (red); Y chromosomes are labeled with retrotransposon *zanzibar* (green). Chromosomes are counterstained with DAPI (blue). Scale bar – 5 μm.

### (c) Meiotic and premeiotic failures in F1 males of interspecies hybrids

To determine possible cytogenetic mechanisms of HMS, we analyzed chromosome behavior in normal-like testes of hybrids from the ♀*An. merus* × ♂*An. coluzzii/An. gambiae* crosses (Figure 2). In primary spermatocytes, homoeologous autosomes pair and form chiasmata in prophase I as in pure species. Unlike pure species, X and Y chromosomes do not pair or display chiasmata in diplotene/diakinesis of prophase I in these hybrids. We found this pattern consistent in all analyzed hybrid males. Metaphase chromosomes in hybrids are visibly longer than at the same stage in pure species, indicating insufficient chromatin condensation. Besides, homoeologous chromosomes in hybrids do not segregate during anaphase. Instead, sister chromatids move to opposite poles of the dividing cell. Because reductional division does not occur in hybrid males, both X and Y chromatids move to the same pole during anaphase. As a result, haploid secondary spermatocytes are not present in these males. Our FISH analysis demonstrated that spermatids in testes of pure species normally contain either an X or Y chromosome. In contrast, we found both X and Y chromosomes present in spermatids of the hybrids (supplementary material, figure S5). Moreover, the abnormal spermatids are larger in size due to insufficient chromatin condensation and the double chromosome content. Thus, we discovered that chromosomes in normal-like testes of hybrid males start with meiotic behavior in prophase and then prematurely switch to mitotic behavior in anaphase, thus, skipping the reductional division. The equational division of primary spermatocytes results in dysfunctional diploid sperm in hybrids when *An. merus* is the mother. Degenerate testes of F1s from the reciprocal ♀*An. coluzzii/gambiae* × ♂*An. merus* crosses have only undifferentiated round germline stem cells. To visualize sex chromosomes, we performed whole-mount FISH with labeled 18S rDNA (Cy3) and satellite AgY53B (Cy5) to mark the sex chromosomes of the hybrid (supplementary material, figure S6). Only interphase sex chromosomes in the nuclei of germline stem cells are detected indicating that meiosis does not start in the underdeveloped testes of F1 hybrids if *An. merus* is the father. Thus, premeiotic arrest in degenerate testes is the reason for the lack of spermatids in these hybrids.

**Figure 2.**
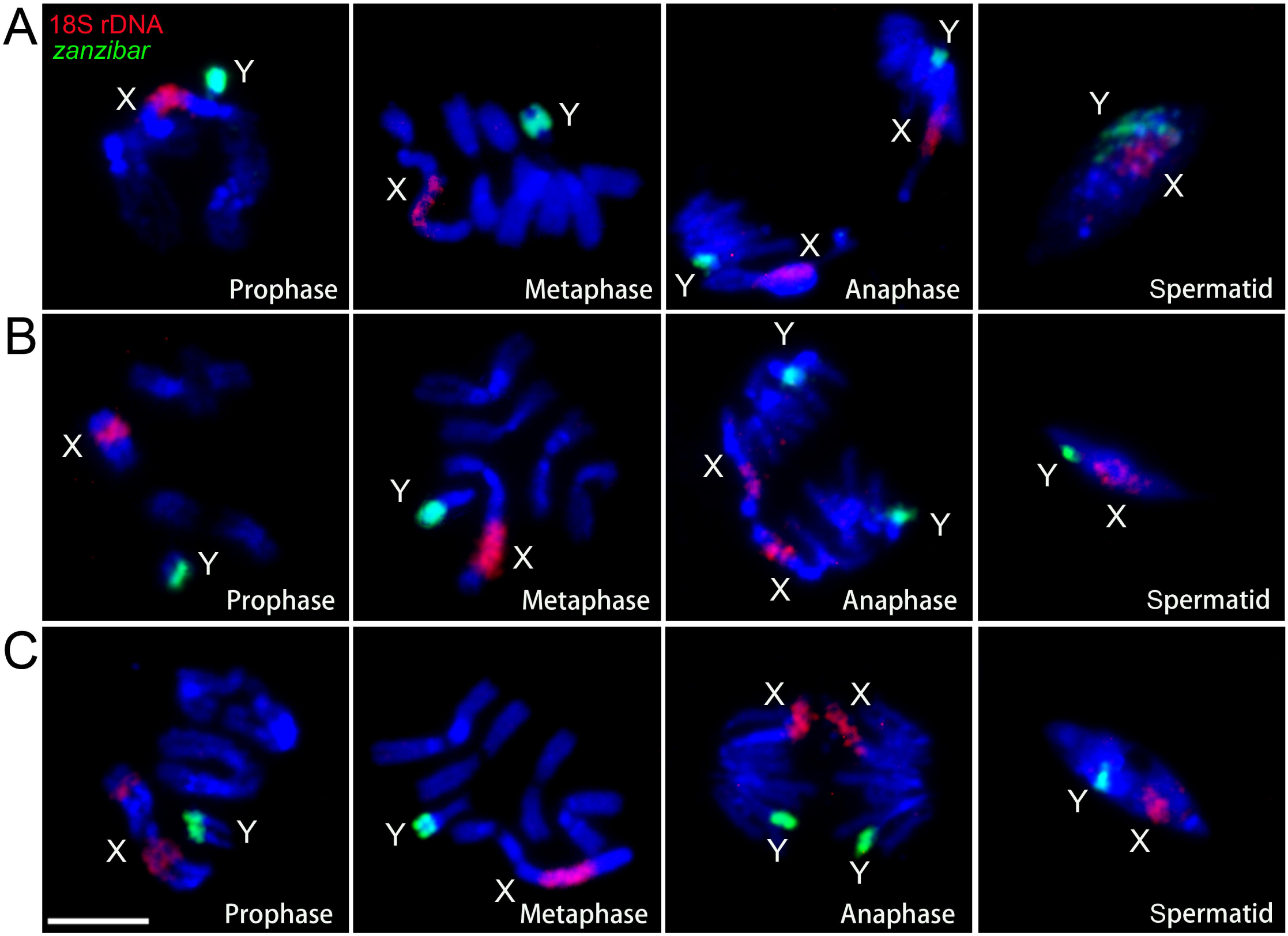
Chromosome behavior during meiosis in testes of interspecies hybrids. (A) ♀*An. merus*♂*An. coluzzii* MALI. (B) ♀*An. merus* × ♂*An. coluzzii* MOPTI. (C) ♀*An. merus* × ♂*An. gambiae* ZANU. X chromosomes are labeled with 18S rDNA (red) and Y chromosomes are labeled with retrotransposon *zanzibar* (green). Chromosomes are counterstained with DAPI (blue). Scale bar – 5 μm.

### (d) Gene expression in pure species and interspecies hybrids

To test if the aforementioned premeiotic or meiotic failures are associated with gene misexpression, we analyzed the transcript profile of the premeiotic and meiotic *vasa* [27], *SMC2*, and *SMC4* genes [28, 29], as well as the meiotic and postmeiotic *Spo11, Msh4, SMC3*β [28, 29], β*2-tubulin* [30, 31], *Ams, mts*, and *Dzip1l* [32] genes in F1 hybrid males (supplementary material, table S1). The RT-PCR results show that all these genes express in the reproductive tissues of *An. coluzzii, An. merus*, and F1 hybrids from the ♀*An. merus* × ♂*An. coluzzii* cross. A slight but noticeable increase was observed for the *vasa, SMC2, SMC4*, and *SMC3*β transcripts in normal-like testes of these hybrids compared with testes of any of the parental species (Figure 3A). This expression profile was consistent between two independent RT-PCR experiments that involved testes combined with MAGs and testes without MAGs. A separate experiment using MAGs only shows no *vasa, SMC2, SMC4*, or *SMC3*β transcripts in this tissue. Although *vasa, SMC2*, and *SMC4* express in degenerate gonads of ♀*An. coluzzii* MOPTI × ♂*An. merus* F1 hybrids, transcripts of all tested meiotic and postmeiotic genes are virtually absent in these hybrids (Figure 3A) supporting our observation that meiosis does not occur in degenerate testes (supplementary material, figure S6). Expression of *vasa* indicated the development of germline stem cells even in degenerate testes of these interspecies hybrids. To test whether gene misexpression occurs in fertile female hybrids, we performed RT-PCR for the same genes using ovaries of pure species and hybrid females (supplementary material, figure S7). Our results show that, unlike males, all analyzed genes have similar transcript profiles in females of pure species and hybrids from the reciprocal crosses.

**Figure 3.**
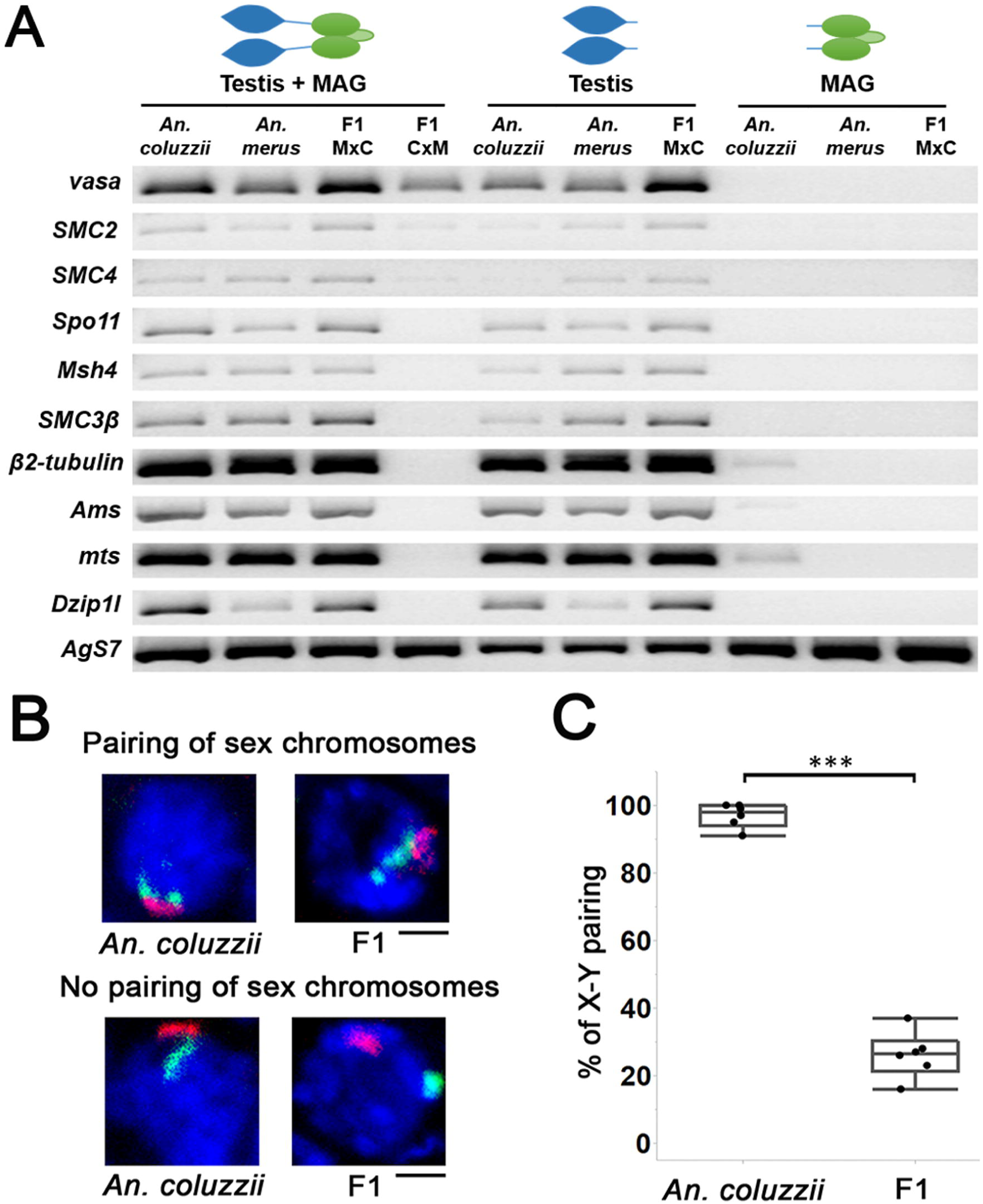
Molecular and chromosomal abnormalities in interspecies hybrids. (A) Gene expression in male reproductive organs of *An. coluzzii* MOPTI, *An. merus*, F1 hybrids of ♀*An. merus* × ♂*An. coluzzii* MOPTI (F1 M×C), and F1 hybrids of ♀*An. coluzzii* MOPTI × ♂*An. merus* (F1C×M) analyzed by RT-PCR. *AgS7* – an endogenous control gene. (B) Confocal images of primary spermatocytes with FISH showing pairing and unpairing of X (red) and Y(green) chromosomes in *An. coluzzii* MOPTI males and F1 hybrid males from the ♀*An. merus* ×♂*An. coluzzii* MOPTI cross. (C) Percentage of the X and Y chromosome pairing in primary spermatocytes of six *An. coluzzii* MOPTI males and six F1 hybrid males from the ♀*An. merus* ×♂*An. coluzzii* MOPTI cross. Statistical significance was assessed with a two-sample pooled *t*-test.

### (e) Chromosomal abnormalities in interspecies hybrids

Here, we performed quantitative analyses of chromatin condensation and X-Y chromosome pairing in pure species and their hybrids. To determine the extent of chromatin condensation in normal-like testes of interspecies hybrids compared with pure species, we measured the lengths of metaphase chromosomes (supplementary material, table S2). The results of statistical analysis with a two-sample pooled *t*-test show that chromosomes in F1 hybrids are typically longer than chromosomes in *An. coluzzii, An. gambiae*, or *An. merus* (supplementary material, figure S8). For example, chromosomes of the *An. merus* origin in a hybrid background always show significant (P<0.001) elongation by at least 1.3 times compared to pure *An. merus*. The X chromosome of *An. merus* suffered the most serious undercondensation in hybrids, exceeding the length of the X chromosome in the pure species background by 1.6-1.9-fold.

In our cytogenetic study of interspecies hybrids from the crosses ♀*An. merus* × ♂*An. coluzzii/An. gambiae*, the X and Y chromosomes do not show pairing or chiasmata in diplotene/diakinesis of prophase I (Figure 2). We hypothesized that sex chromosome pairing is affected in early prophase I when individual chromosomes cannot be distinguished by direct visualization. To analyze the X-Y chromosome pairing at the pachytene stage of prophase I, we performed a whole-mount FISH and examined spatial positions of the X- and Y-specific fluorescent signals in confocal optical sections of nuclei in testes of pure species and their hybrids (supplementary material, figure S9). We recorded the number of nuclei with X and Y fluorescent signals colocalized versus X and Y fluorescent signals located separately (Figure 3B, supplementary material, table S3). The results of a statistical analysis with a two-sample pooled *t*-test demonstrate that X and Y chromosomes pair in more than 90% of primary spermatocytes in *An. coluzzii*, while they pair in less than 30% of primary spermatocytes in F1 hybrids of ♀*An. merus* ♂*An. coluzzii* (Figure 3C).

## 4. Discussion

In this study, we performed the first detailed cytological analysis of spermatogenesis in pure species and hybrids of mosquitoes. We demonstrate that premeiotic and meiotic defects are involved in HMS in reciprocal crosses (Figure S10). This observation is at odds with the commonly accepted view that hybrid males suffer postmeiotic sterility problems more often than premeiotic or meiotic sterility problems [1, 33]. Although many postmeiotic defects seen in *Drosophila* hybrids are indeed related to problems in sperm bundling and motility [5, 9, 33], some sperm abnormalities may stem from meiotic failures. For example, our data shows that sperm abnormalities such as nonmotility, two-tailed heads, and chromatin decompaction in sperm heads can result from impaired meiosis I (Figure 2). Additional studies of interspecies hybrids of various organisms should determine whether the first meiotic division commonly fails when fertility is affected.

Our cytogenetic investigation of meiosis in *Anopheles* species demonstrates that the X and Y chromosomes pair with each other throughout prophase I (Figure 1, 3BC, supplementary material, figures S3, S4). Unlike pure species, meiotic prophase I in F1 mosquito hybrids shows cytogenetic anomalies—low percentages of X-Y chromosome pairing and insufficient chromatin condensation (Figure 3BC, supplementary material, figure S8). The interplay between these two phenotypes may result in failing a reductional meiotic division and prematurely proceeding to an equational mitotic division (Figure 4). All chromosomes must achieve synapsis in pachynema, complete DNA repair, and disassemble their synaptonemal complexes in diplonema as homologous chromosomes prepare to segregate [34]. The presence of chiasmata and tension exerted across homologs ensures that cells undergo reductional segregation [35]. Asynapsis of chromosomes almost invariably triggers pachytene checkpoint and meiotic breakdown [36].

**Figure 4.**
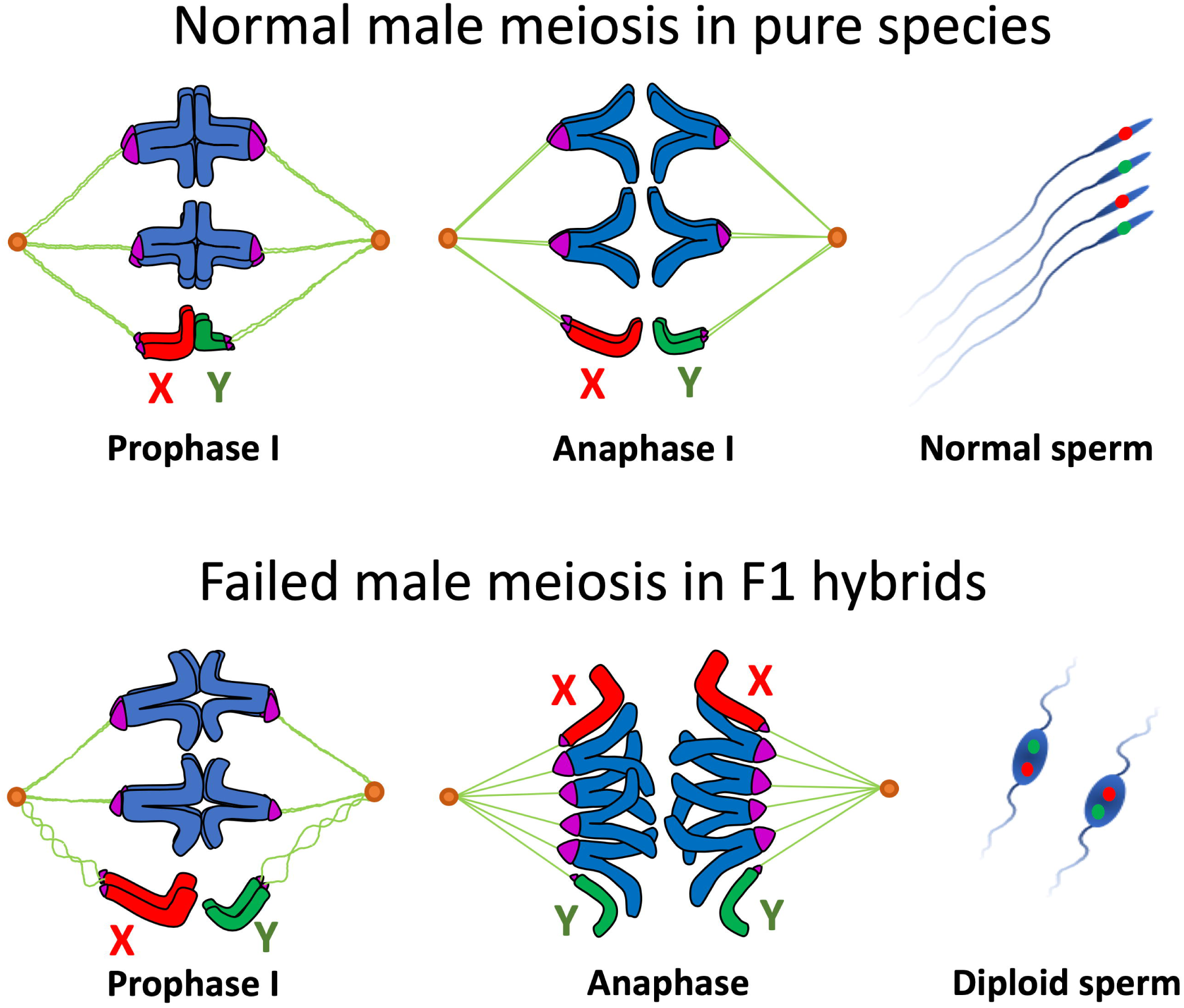
Scheme of male meiosis in pure species and interspecies hybrids of the *An. gambiae* complex. Unlike pure species, X and Y chromosomes in meiotic prophase I of hybrids are largely unpaired. The absence of chiasmata between the sex chromosomes and insufficient chromatin condensation reduce tension exerted across the homoeologous chromosomes, triggering premature sister chromatid segregation. As a result, instead of a reductional division, primary spermatocytes in interspecies hybrids undergo an equational mitotic division leading to diploid nonmotile sperm.

Haldane’s rule predicts that meiosis would be aberrant in males but may be normal in females [20]. This could be due to differences in the mechanisms of X-Y and X-X pairing. For example, in *D. melanogaster*, X and Y chromosomes pair through specific pairing sites located in heterochromatin [37], which could be relatively quickly altered during evolution. In contrast, the X-X and autosome-autosome pairing capacity is widely distributed along the chromosome arm[38] and is likely more robust to evolutionary changes. A study of introgression of the Y chromosome from *An. gambiae* into *An. arabiensis* suggests that the Y chromosome does not cause sterility [17]. This runs counter to our findings that X-Y unpairing underlies sterility in male hybrids. Identification and analysis of X-Y pairing sites in species of the *An. gambiae* complex may shed light on this problem.

Reciprocal crosses in malaria mosquitoes and other organisms usually produce hybrids with sterility phenotypes of different degrees of severity [6, 18, 39]. This observation indicates that malfunction of multiple stages of spermatogenesis can be involved in postzygotic isolation. At the early stages of speciation, premeiotic, meiotic, and postmeiotic genes still express in hybrids and meiotic errors are characterized by decreased pairing between sex chromosomes, insufficient chromatin condensation, and skipping reductional division. These phenotypes are seen in sterile hybrids from the ♀*An. merus* × ♂*An. coluzzii/An. gambiae* crosses (Figures 2, 3, 4). Higher expression levels of the *vasa, SMC2*, and *SMC4* genes (Figure 3A) are consistent with our observation of extended premeiotic and meiotic stages in these hybrids (Figure S2). It is possible that the observed overexpression of *SMC3*β in the hybrid males may contribute to abandoning homologous chromosome segregation in favor of sister-chromatid cohesion, recombination, and premature segregation. This is because the SMC1 and SMC3 proteins function together with other proteins to control cell cycle transitions by participating in sister-chromatid cohesion, recombination, and faithful segregation [29, 40, 41]. To support crossover formation between homologous chromosomes and their subsequent segregation in meiosis, repair from the more readily available homologous sequences on the sister chromatid must be suppressed [34].

As species continue to diverge, meiotic errors become more prominent, and new hybrid phenotypes appear as has been seen, for example, in sterile male hybrids from the ♀*D. pseudoobscura* × ♂*D. persimilis* cross [6, 7]. At this stage of postzygotic isolation, meiosis is manifested by the unpairing of most of the chromosomes and by malfunction of the abnormally elongated spindle, resulting in spermatids with unbalanced chromosome content in sterile male hybrids [6, 7]. In mice hybrids, spermatogenic arrest occurs in early meiosis I when homoeologous chromosomes fail to pair and meiotic sex chromosome inactivation is disrupted, resulting in increased apoptosis of spermatocytes [3, 4, 42]. Disruption of synapsis between heterospecific chromosomes in prophase I has been proposed as a recurrently evolving trigger for the meiotic arrest of interspecific F1 hybrids [3, 43]. Finally, spermatogenic abnormalities in hybrids can happen before meiosis starts. A spermatogenic arrest at the premeiotic stage is characterized by the repression of meiotic and postmeiotic genes and by the lack of spermatocytes in degenerate testes of hybrid males. These phenotypes have been observed in sterile hybrids from the ♀*D. mauritiana* × ♂*D. sechellia* cross [5] and in sterile hybrids from the ♀*An. coluzzii/An. gambiae* × ♂*An. merus* crosses (our study).

Charles Darwin, in the chapter on “Hybridism” in his book, the *Origin of Species*, rightly argued that hybrid sterility ‘‘is not a specially endowed quality, but is incidental on other acquired differences [44].’’ Identification of the germ-line-specific cellular and molecular differences acquired during the early stages of postzygotic isolation between species is crucial to explaining both speciation and normal fertility function. Our study suggests that meiotic abnormalities in hybrid males stem from the unpairing of the sex chromosomes and insufficient chromatin condensation. Cytological analyses of spermatogenesis in interspecies hybrids from diverse groups of organisms may highlight general patterns and mechanisms in the origin and evolution of postzygotic isolation. Recently evolved members of the *An. gambiae* complex represent an excellent, new system for studying the genetic basis and molecular mechanisms of species incompatibilities at the early stages of postzygotic reproductive isolation.

## Supporting information

Five DNA probes were used for FISH in this study (supplementary material, table S1)

the ruler tool in Adobe Photoshop CS6 (Adobe Inc., San Jose, CA, USA) (supplementary material, table S2)

were analyzed in pure species and hybrids, respectively (supplementary material, table S3)

vibrant motility escaped from the ruptures (supplementary material, figure S1A, movie S1)

two short tails growing from opposite ends of the head (supplementary material, figure S1B, movie S2)

## Acknowledgments

The following mosquito strains were obtained through BEI Resources, NIAID, NIH: *An. coluzzii*, Strain Mali-NIH, Eggs, MRA-860 contributed by Nora J. Besansky; *An. coluzzii*, Strain MOPTI, Eggs, MRA-763 contributed by Gregory C. Lanzaro; *An. gambiae*, Strain ZANU, MRA-594 contributed by Hilary Ranson and Frank H. Collins; *An. merus*, Strain MAF, MRA-1156 contributed by Maureen Coetzee. We thank Kristin Rose and Janet Webster for editing the text.

