## Supplementary material for "Premeiotic and meiotic failures lead to hybrid male sterility in the *Anopheles gambiae* complex": Five DNA probes were used for FISH in this study (supplementary material, table S1)

### Supplemental Material

This file contains

- Supplementary Methods
- Supplementary Table S1
- Supplementary Figures S1-10
- Supplementary References

#### Supplementary Methods

##### Mosquito strains and crossing experiments

The laboratory colonies of *An. gambiae* ZANU (MRA-594), *An. coluzzii* MOPTI (MRA-763), *An. coluzzii* MALI (MRA-860), and *An. merus* MAF (MRA-1156) were obtained from the Biodefense and Emerging Infections Research Resources Repository (BEI). Authentication of the species was performed by a cytogenetic analysis and by PCR diagnostics [1, 2]. Mosquitoes were reared at 27±1°C, with a 12-hour photoperiod and 70±5% relative humidity. Larvae were fed fish food, and adult mosquitoes were fed 1% sugar water. To induce oviposition, females were fed defibrinated sheep blood (Colorado Serum Co., Denver, Colorado, USA) using artificial blood feeders. To perform interspecies crosses, male and female pupae were separated to guarantee virginity of adult mosquitoes. We differentiated males and females at the pupal stage using sex-specific differences in the shape of their terminalia [3]. After the emergence of adults, crossing experiments were performed by combining 30 females and 15 males in one cage. Five days after random mating, the females were fed sheeps blood.  Two days later, an egg dish, covered with moist filter paper to keep the eggs from drying out, was put into the cage. Backcrosses of F1 males and parental females were done using a similar method. At least two blood meals were fed to females and at least three repeats of each cross were conducted.

**Male gonad and sperm observation**

Male gonads were dissected using an Olympus SZ61 stereo microscope (Olympus, Tokyo, Japan) and photographed with an Olympus BX41 phase contrast microscope and a UC90 digital camera (Olympus, Tokyo, Japan). We observed the testes of 30 males and took pictures of the testes of five males from each cross. For sperm observation, testes were mounted in 20 µl sperm assay buffer containing 4 mM KCl, 1.3 mM $CaCl_2$, 145 mM NaCl, 5 mM D-glucose, 1 mM $MgCl_2$, and 10 mM 4-(2-hydroxyethyl)-1-piperazineethanesulfonic acid (Hepes) [4]. After gently covering testes with a coverslip, they were crushed, and sperm motility was observed under a BX41 phase contrast microscope (Olympus, Tokyo, Japan). A movie of sperm motility for at least five hybrid males from each cross and five pure species males was recorded using a UC90 digital camera (Olympus, Tokyo, Japan).

**Chromosome preparation**

Testes with male accessory glands were dissected from male pupae and 0-12 hours-old adults placed in 0.075% potassium chloride (KCl) hypotonic solution on a frosted glass slide (Thermo Fisher Scientific, Waltham, MA, USA) under an Olympus SZ61 dissecting microscope (Olympus, Tokyo, Japan). To observe meiotic chromosomes, male accessory glands and other tissues were removed, and only testes were left on the slide. Immediately after dissection, a drop of 50% propionic acid was added to the testes, and they were covered with a 22×22-mm coverslip (Thermo Fisher Scientific, Waltham, MA, USA). Preparations were tapped using the flat rubber end of a pencil and observed under an Olympus CX41 phase-contrast microscope (Olympus, Tokyo, Japan). The slides were frozen in liquid nitrogen and a sharp razor was used to take off the coverslips. The preparations were placed in 50% ethanol chilled at -20ºC for a minimum of 2 hours. Later, serial dehydrations were performed in 70%, 90%, and 100% ethanol for 5 min each at room temperature (RT). Subsequently, the best preparations from at least 10 slides/individuals with desired meiotic stages were chosen for further studies.

**DNA probe labeling**

Five DNA probes were used for fluorescence *in situ* hybridization (FISH) in this study (Table S1): retroelement *Zanzibar*, which is specific to the Y chromosome of *An. gambiae* and *An. coluzzii*, 18S rDNA, which is specific to the X chromosome of *An. gambiae* and *An. coluzzii* but hybridized to both the X and Y chromosomes of *An. merus,* satellite AgY53B, which is derived from the AgY53B/AgY477 satellite array and hybridized to X and Y chromosomes of all three species [5], satellite AgY477-AgY53B junction region, which is specific to the Y chromosome of *An. gambiae* and *An. coluzzii*, and a satellite from Contig_240 identified from *An. gambiae* sequencing data (SRS667972, SRR1509742, SRR1508169) [5] using the Redkmer pipeline [6], which is specific to the X chromosomes of *An. coluzzii* and *An.  merus*. The 18S rDNA, AgY53B, and *zanzibar* probes were labeled by the Cyanine3 (Cy3) or Cyanine5 (Cy5) fluorochromes using PCR with genomic DNA as a template. Genomic DNA was extracted using DNeasy Blood & Tissue Kits (Qiagen, Hilden, Germany) from virgin males of *An. gambiae* ZANU or *An. coluzzii* MOPTI for labeling *zanzibar*, virgin males of *An. merus* MAF for labeling satellite AgY53B, and virgin females of *An. gambiae* or *An. coluzzii* for labeling 18S rDNA.

Each 25 μl of a PCR mix consisted of 1-μl genomic DNA, 12.5-μl ImmoMix™ 2× reaction mix (Bioline USA Inc., Taunton, MA, USA), 1 μl of 10-μM forward and reverse primers, and water. PCR labeling was performed in the Mastercycler® pro PCR thermocycler (Eppendorf, Hamburg, Germany) starting with a 95°C incubation for 10 min followed by 35 cycles of 95°C for 30 sec, 52°C for 30 sec, 72°C for 45 sec, 72°C for 5 min, and a final hold at 4°C. Oligonucleotides with 3'-end- labeling by Cy5 or Cy3 fluorochromes (Sigma-Aldrich, St. Louis, MO, USA) were designed based on sequences of the satellite AgY477-AgY53B junction region, and a satellite from Contig_240.

**FISH**

FISH was performed as previously described [7, 8]. Briefly, slides with good preparations were treated with 0.1-mg/ml RNase at 37°C for 30 min. After washing with 2×saline-sodium citrate (SSC) for 5 min twice, slides were digested with 0.01% pepsin and 0.037% HCl solution for 5 min at 37°C. After washing slides in 1×phosphate-buffered saline (PBS) for 5 min at RT two times, preparations were fixed in 3.7% formaldehyde for 10 min at RT. Slides were then washed in 1×PBS and dehydrated in a series of 70%, 80%, and 100% ethanol for 5 min at RT. Then, 10 μl of probes were mixed, added to the preparations, and incubated at 37°C overnight. After washing slides in 1×SSC at 60°C for 5 min, 4×SSC/NP40 solution at 37°C for 10 min, and 1×PBS for 5 min at RT, preparations were counterstained with a DAPI-antifade solution (Life Technologies, Carlsbad, CA, USA) and kept in the dark for at least 2 hours before visualization with a fluorescent microscope.

**Cytogenetic analyses**

Cytogenetic analyses of FISH-labeled chromosomes were performed on at least 10 male individuals of each pure species and of each hybrid. To visualize and photograph meiotic chromosomes after FISH, we used an Olympus BX41 microscope (Olympus, Tokyo, Japan) with a connected UC90 digital camera (Olympus, Tokyo, Japan). Because retroelement *zanzibar* is absent in *An. merus*, we used satellite AgY53B and 18S rDNA to label sex chromosomes in this species. Each of these probes hybridized with both X and Y chromosomes in *An. merus*, making discrimination between X and Y more difficult in this species. However, we could differentiate metaphase sex chromosomes by the euchromatic arm of the X chromosome and by the slightly larger distal heterochromatic block of the Y chromosome (Figure S4). Heterochromatic parts of the X and Y chromosomes are relatively large and structurally similar in *An. merus* compared to *An. gambiae* or *An. coluzzii*. In the latter two species, the heterochromatic parts of the X and Y chromosomes are relatively small and substantially different from each other both in size and genetic content [5]. Lengths of well-spread metaphase I chromosomes of each species and of each hybrid were measured using the ruler tool in Adobe Photoshop CS6 (Adobe Inc., San Jose, CA, USA) (Table S2). Since the variances for both groups were equal ($s_{max}/s_{min} < 2$), a statistical two-sample pooled *t*-test was done with JMP 13 software (SAS Institute Inc., Cary, NC, USA).

**Whole-mount FISH and analysis of chromosome pairing**

Testes from one-day-old adults of pure species and hybrids were dissected in 1×PBS solution and fixed in 3.7% paraformaldehyde in 1×PBS with 0.1% tween-20 (PBST) for 10 min at room temperature. After incubation with 0.1-mg/ml RNase for 30 min at 37 ºC, testes were penetrated with 1% triton/0.1M HCl in PBST for 10 min at room temperature. The *An. merus* X chromosomes were labeled with 18S rDNA (Cy3) and a satellite from Contig_240 (Cy3), while the *An. coluzzii* MOPTI Y chromosomes were labeled with satellite AgY53B (Cy5) and the satellite AgY477-AgY53B junction region (Cy5). After adding labeled DNA probes, testes were incubated at 75 ºC for 5 min (denaturation) and 37 ºC overnight (hybridization). Later, testes were washed with 2×SSC and mounted with a DAPI-antifade solution (Life Technologies, Carlsbad, CA, USA). Testes from 6 individuals of pure species and of hybrids were scanned and analyzed. Visualization and z-tack 3D scanning were performed on the whole testes with an interval of 1.25 µm between two optical sections under a 63× oil lens of a Zeiss LSM 880 confocal laser scanning microscope (Carl Zeiss AG, Oberkochen, Germany). Based on the size of the testis cells and on the limitations of microscope imaging, we chose to analyze sex-chromosome-pairing events from the first to sixteenth optical layers with strong and clearly detected fluorescent signals. Specifically, cell nuclei at the early stages of meiotic prophase I from the $4^{th}\pm1$, $8^{th}\pm1$, and $12^{th}\pm1$ optical layers were used to count paring events. A total of 418 and 489 nuclei at the early stages of meiotic prophase I were analyzed for pairing and unpairing of the sex chromosomes in pure species and hybrids, respectively (Table S3). The percentage of the number of cells with pairing and no pairing of sex chromosomes was used to compare parental species and hybrids. Since the variances for both groups were equal ($s_{max}/s_{min} < 2$), a statistical two-sample pooled *t*-test was done using JMP 13 software (SAS Institute Inc., Cary, NC, USA).

**RNA extraction and RT-PCR**

We dissected reproductive organs, which include both testes and male accessary glands (MAGs), from 30 males, only testes from 20 males, only MAGs from 20 males, and ovaries from 30 females. These mosquitoes were 0-12-hour-old virgin adults of *An. coluzzii* MOPTI, *An. merus* MAF, and interspecies hybrids from the ♀*An. coluzzii* MOPTI × ♂*An. merus* and ♀*An. merus* × ♂*An. coluzzii* MOPTI crosses. Total RNA was extracted using a Direct-Zol™ RNA MiniPrep Kit (Zymo Research, Irvine, California, US). Because testes and MAGs in F1 hybrids from ♀*An. coluzzii* MOPTI × ♂*An. merus* crosses are underdeveloped, they are too difficult to dissect. Therefore, we did not separate testes from MAGs. Total RNA was diluted in 50 µl diethyl pyrocarbonate (DEPC)-treated water and the concentration was measured by NanoDrop™ 2000 Spectrophotometers (Thermo Fisher Scientific, Waltham, MA, USA). Two-step RT-PCR was performed to analyze gene expression. cDNA for selected genes (Table S1) was generated using a SuperScript™ III First-Strand Synthesis System for RT-PCR (Invitrogen, Carlsbad, CA, US). For the protocol, a 10 µl mixture, including 2 µl of total RNA, 1 µl of 50 µM oligo$(dT)_{20}$ primer, 1 µl of 10 mM dNTP mix solution, and 6 µl of DEPC-treated water, was incubated at 45°C for 5 min. After being chilled on ice for at least 1 min, 10 µl of cDNA synthesis mix containing 2 µl of 10× RT buffer, 1 µl of 25mM $MgCl_2$, 2 µl of 0.1-M dithiothreitol (DTT), 1 µl of RNaseOUT™ (40 U/µl), and 1 µl of SuperScript™ III RT (200 U/µl) was added into each RNA/primer mixture. After incubation at 50°C for 50min, reactions were terminated at 85°C for 5 min. Later, 1 µl of RNase H was added into each tube for a 20 min incubation at 37°C. Next, the cDNA synthesis reaction was used for PCR. Each 25-µl PCR mix consisted of 1-µl cDNA, 12.5-µl ImmoMix™ 2× reaction-mix (Bioline USA Inc., Taunton, MA, USA), 1 µl of 10-µM forward and reverse primers, and water. PCR was performed in the C1000 Touch™ thermal cycler platform (Bio-Rad Laboratories, Hercules, CA, US) starting with a 10 min incubation at 95°C followed by 32 cycles of 95°C for 30 sec, 57°C for 30 sec, 72°C for 30 sec, 72°C for 10 min, and a final hold at 4°C. Amplification products were visualized in a 2% agarose gel and photographed under the same parameters for each gene. Each RT-PCR experiment was repeated three times with each sample to ensure consistency of the results.

#### Supplementary Table

Table S1. Labeled oligos and primer sequences used for FISH and RT-PCR.

| **Sequence name** | **GenBank accession number** | **VectorBase gene ID** | **Labeled oligos and PCR primer sequences** | **Reference to oligo design** |
|---|---|---|---|---|
| **18S rDNA** | AM157179 | AGAP028978 | F: AACTGTGGAAAAGCCAGAGC | This study |
| | | | R: TCCACTTGATCCTTGCAAAA | |
| ***zanzibar*** | KP878482 | | F: TTCTTCGATGTTGTGCTGGA | [5] |
| | | | R: ATGGAGAAACAGGGCAACAA | |
| | | | F: ATGCATGCTTGGATTCCTTC | |
| | | | R: GGTTTCTATGATCGCCTGGA | |
| | | | F: TTGGCATTCATCTGTCCAAA | |
| | | | R: GCACCCTTGATCTCATGTCA | |
| **AgY53B** | AY754156 | | F: CCTTTAAACACATGCTCAAATT | [9] |
| | | | R: GTTTCTTCATCCTTAAAGCCTAG | |

| | | | | |
|---|---|---|---|---|
| **AgY477-AgY53B junction region** | AY754151 | | TTCTAAGTTTCTAGGCTTTAAGGAT GAAGAAACCGACTATTC[Cyanine5] | This study |
| **Contig_240** | SRS667972, SRR1509742, SRR1508169 | | CAATAAATTTCCTTTTTAATGATGC AAAATCTACGTCTCTAGC[Cyanine3] | This study |
| ***vasa*** | XM_314684 | AGAP008578 | F: TTCTGCTGAGGTGCTTAGCG<br>R: CGTCTCCGCTCATGTTTCCT | This study |
| ***SMC2*** | XM_554796 | AGAP011425 | F: GATCAATGGCAAGTCGGTGC<br>R: CTTTTTGCTTTCCACCGCCA | This study |
| ***SMC4*** | XM_320297 | AGAP007826 | F: AACTCAGCGAAGCATCCGAA<br>R: GCGTCGTACGTTTCATCGTG | This study |
| ***Spo11*** | XM_309793 | AGAP010898 | F: CGATTGCAATGGTCGATGGG<br>R: TCAATCTCCACCTTGGTCGC | This study |
| ***Msh4*** | XM_317674 | AGAP012245 | F: AACGCACCAAAACACGGTTC<br>R: GTATGCTCTCGATCAGCCCC | This study |
| ***SMC3β*** | XM_557814 | AGAP008672 | F: CGGTCCGATAGTGCGTATGT<br>R: TTTCCTTCCACTCTTGCCCG | This study |
| ***β2-tubulin*** | XM_314718 | AGAP008622 | F: GTACGTGCCGGATCATTTCG<br>R: GGCCAGTTTGCAAATGCACTA | This study |
| ***Ams*** | FJ869235 | AGAP029148 | F: CATACGGGAGGTGAGGAAAT<br>R: CCCCTTCATGCTTCATCTT | [10] |
| ***mts*** | FJ869236 | AARA006451 | F: TGGGATCCAAATTATTTCGTG<br>R: CTGTTCGGTTCAACAATGGA | |

| | | | | |
|---|---|---|---|---|
| ***Dzip1l*** | FJ869237 | AGAP001165 | F: GGCCAAAGTGATACAAATTGTTT | |
| | | | R: CGTTTCCAATAGGGACTTCG | |
| ***AgS7*** | L20837 | AGAP010592 | F: AGAACCAGCAGACCACCATC | [11] |
| | | | R: GCTGCAAACTTCGGCTATTC | |

***SMC*** – Structural Maintenance of Chromosomes

***SMC3β*** – paralog of SMC3 (AGAP006388)

***Spo11*** – initiator of meiotic double strand breaks

***Msh4*** – Mutator S homolog 4

***Ams*** – *Anopheles* male specific

***mts*** – mosquito testis specific

***Dzip1l*** – DAZ (deleted in azoospermia) interacting zinc finger protein 1-like

***AgS7*** – *Anopheles gambiae* 40S ribosomal protein S7

#### Supplementary Figures

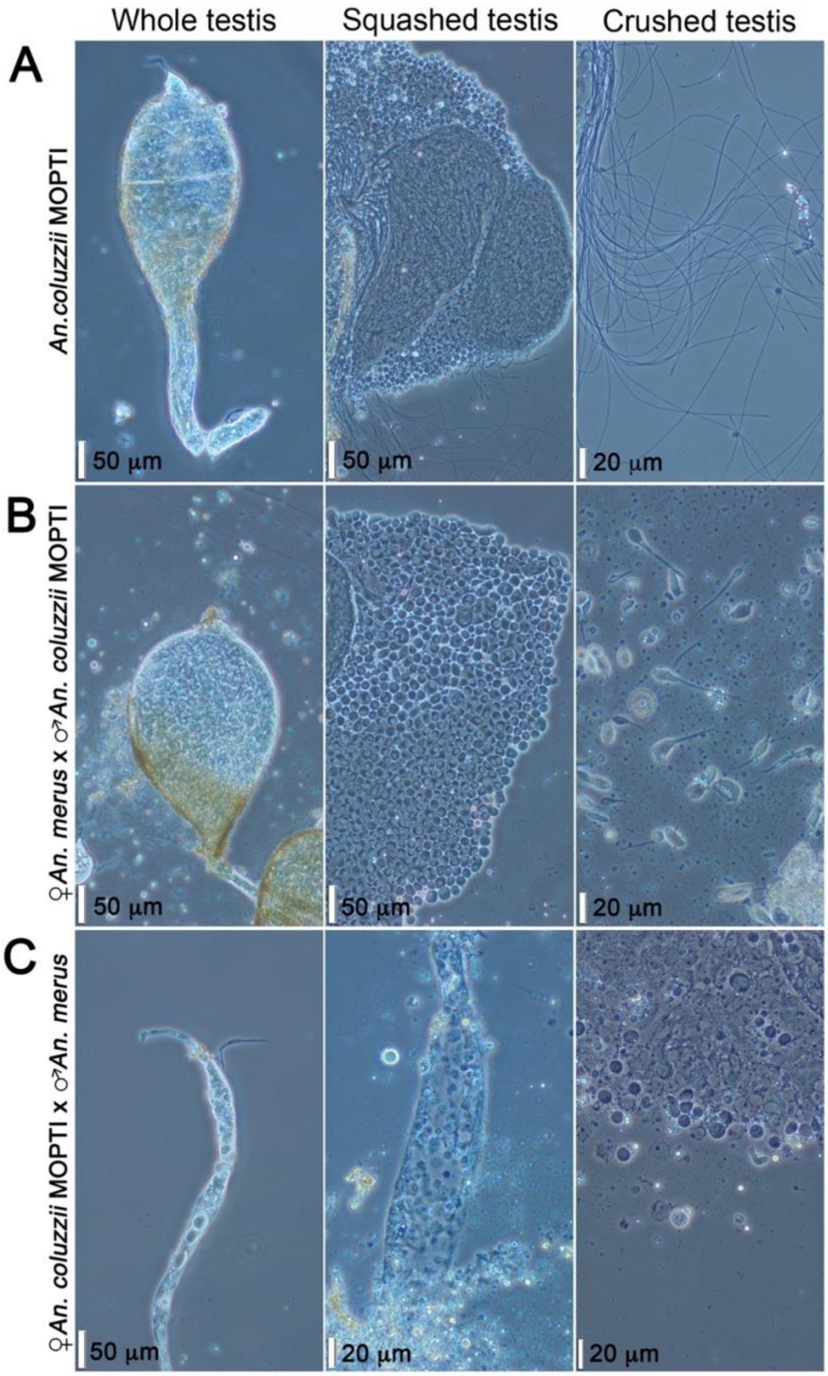


Figure S1. Morphology and cell content of testes from 2-day-old adult males. (A) Whole testis, squashed testis, and mature spermatozoa in crushed testis from *An. coluzzii* MOPTI. (B) Normal-like whole testis, squashed testis, and immature spermatozoa and spermatids in a crushed testis from an F1 hybrid of the ♀*An. merus* × ♂*An. coluzzii* MOPTI cross. (C) Underdeveloped whole testis, squashed testis, and undifferentiated cells in a crushed testis from an F1 hybrid of the ♀*An. coluzzii* MOPTI × ♂*An. merus* cross. Vertical scale bars – 50 and 20 µm.

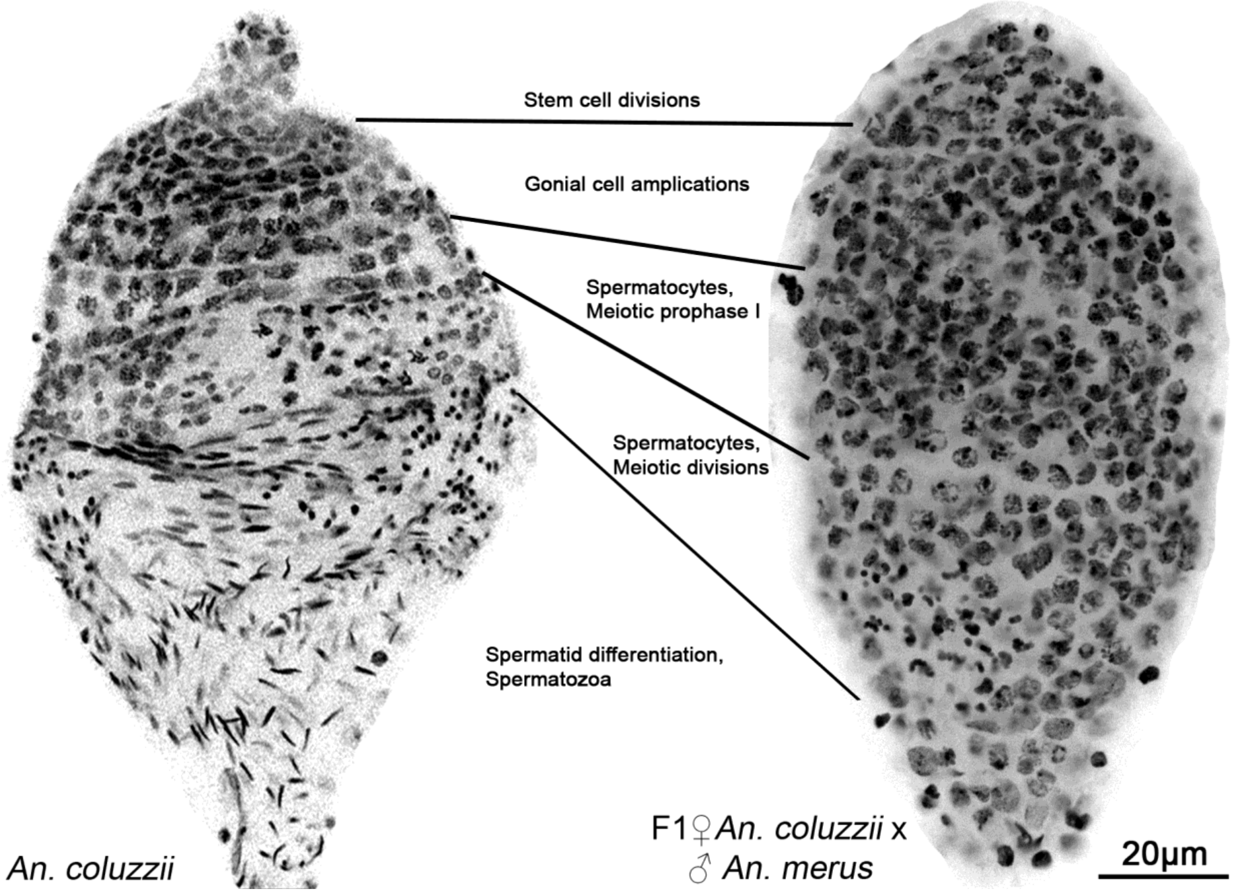


Figure S2. Spermatogenesis comparison between a testis from a one-day-old adult of *An. coluzzii* MOPTI (left) and a normal-like testis from a one-day-old adult of an F1 hybrid of the ♀*An. merus* × ♂*An. coluzzii* MOPTI cross (right). Chromatin is counterstained with DAPI (black). Stages of spermatogenesis are determined by cell morphologies and visual boundaries between the stages.

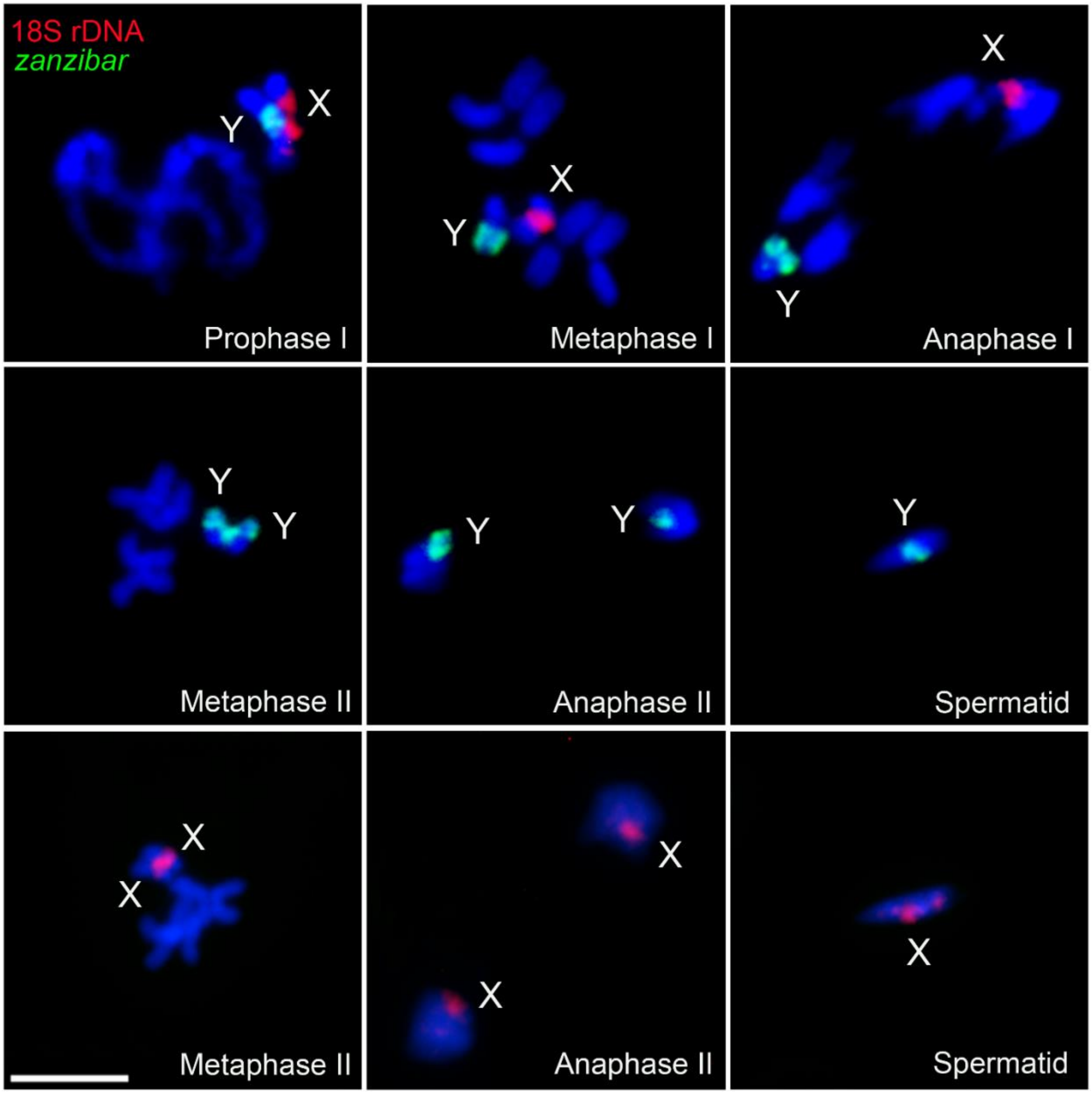


Figure S3. Chromosome behavior during meiosis in testes of *An. coluzzii* MOPTI. X chromosomes are labeled with 18S rDNA (red), and Y chromosomes are labeled with retrotransposon *zanzibar* (green). Chromosomes are counterstained with DAPI (blue). Scale bar – 5 μm.

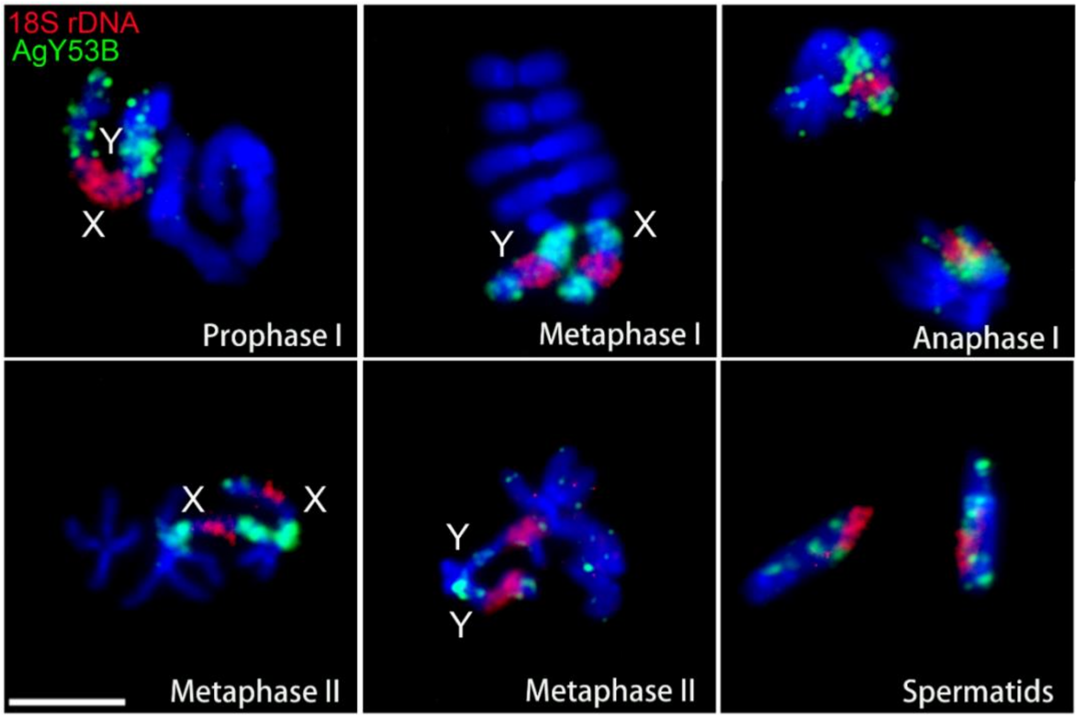


Figure S4. Chromosome behavior during meiosis in testes of *An. merus* MAF. Sex chromosomes are labeled with 18S rDNA (red) and satellite AgY53B (green). Chromosomes are counterstained with DAPI (blue). Scale bar – 5 µm.

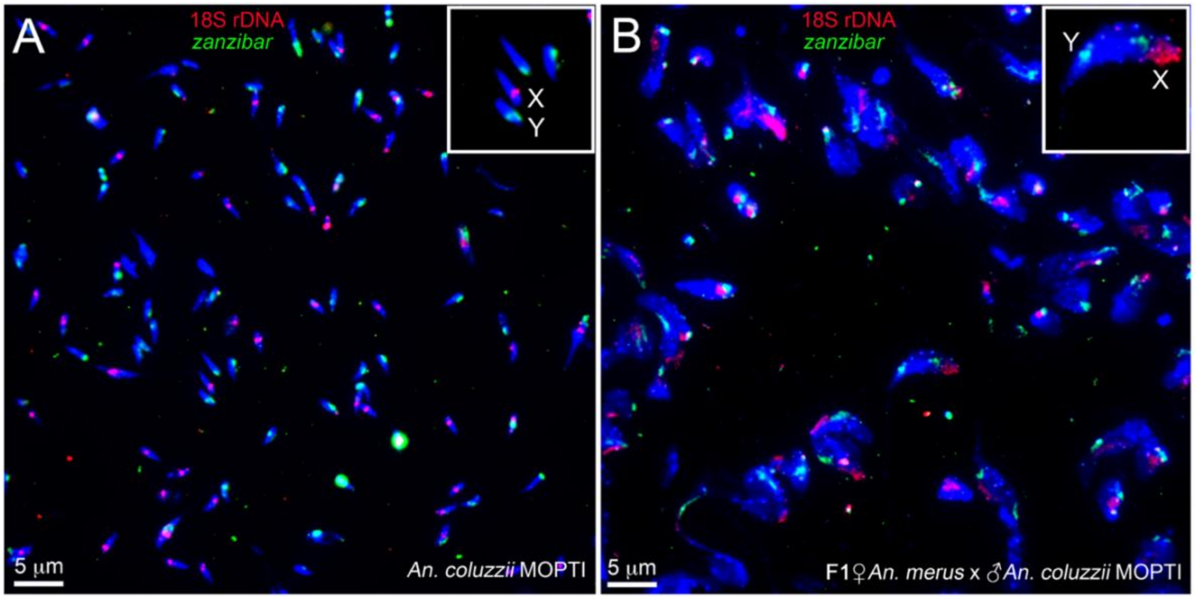


Figure S5. Sizes and sex chromosome content of spermatids in pure species and interspecies hybrids. (A) Spermatids from a 2-day-old adult of *An. coluzzii* MOPTI. (B) Spermatids from a 5-day-old adult F1 hybrid from the ♀*An. merus* × ♂*An. coluzzii* MOPTI cross. The X and Y chromosomes are visualized with 18S rDNA (red) and retrotransposon *zanzibar* (green), respectively. Chromatin is counterstained with DAPI (blue). The insets show magnified images of spermatids.

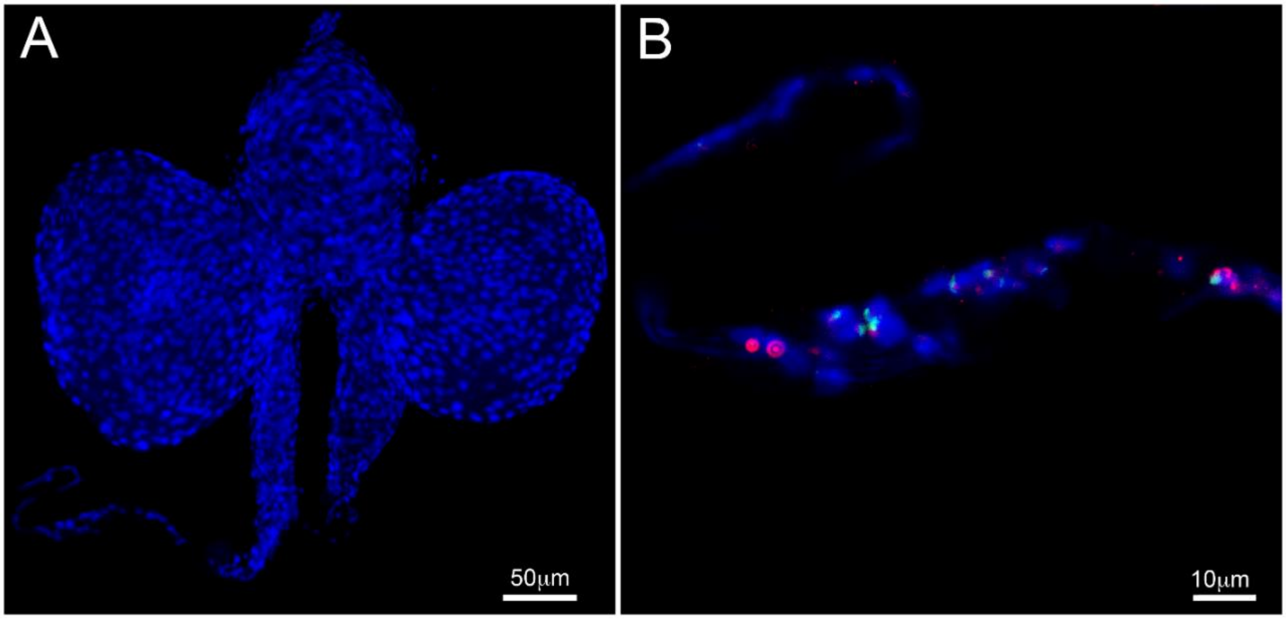


Figure S6. Visualization of sex chromosomes in a degenerate testis of a hybrid adult from the ♀*An. coluzzii* MOPTI × ♂*An. merus* cross. (A) DAPI-stained male accessory glands and an underdeveloped testis of a one-day-old hybrid adult. (B) Whole-mount FISH of the degenerate testis with 18S rDNA (red) and satellite AgY53B (green) detects the interphase sex chromosomes. Chromatin is counterstained with DAPI (blue).

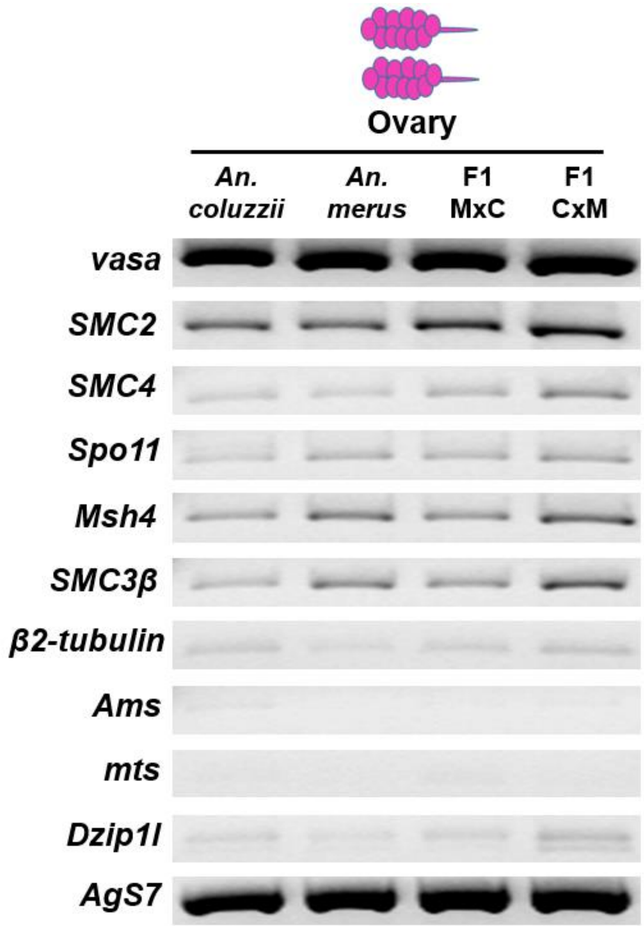


Figure S7. Gene expression in ovaries of *An. coluzzii*, *An. merus*, F1 hybrids of ♀*An. merus* × ♂*An. coluzzii* MOPTI (F1 M×C), and F1 hybrids of ♀*An. coluzzii* MOPTI × ♂*An. merus* (F1C×M) analyzed by RT-PCR. *AgS7* – an endogenous control gene.

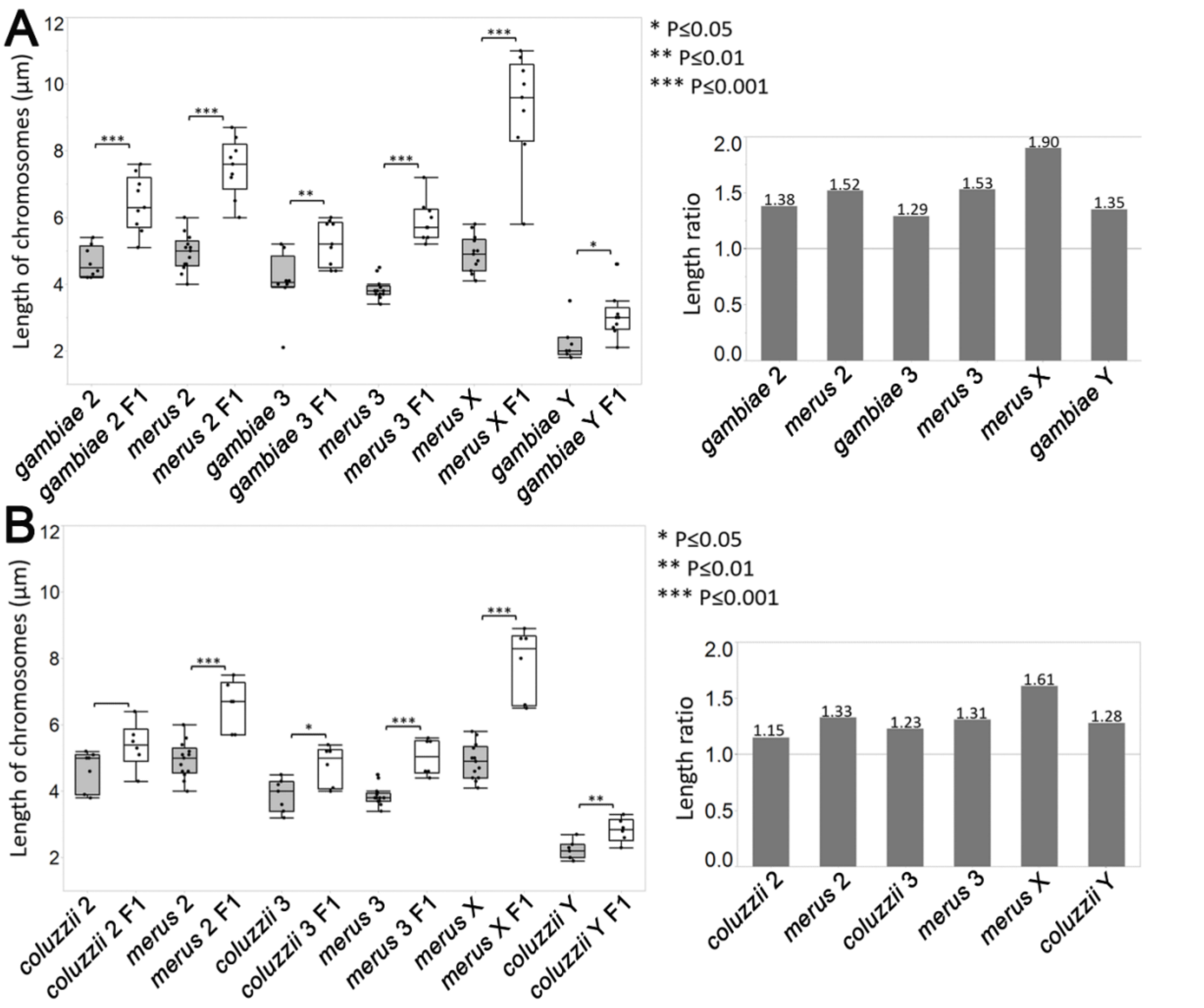


Figure S8. Increased lengths of chromosomes in testes of interspecies F1 hybrids. (A) Left panel: Lengths of metaphase chromosomes in *An. gambiae* ZANU males (filled boxplots) and in F1 hybrid males from the ♀*An. merus* × ♂*An. gambiae* ZANU cross (open boxplots). Right panel: Ratios of the lengths of chromosomes between F1 hybrids and *An. gambiae* ZANU. (B) Left panel: Lengths of metaphase chromosomes in *An. coluzzii* MOPTI (filled boxplots) and in F1 hybrid males from the *An. merus* × ♂*An. coluzzii* MOPTI cross (open boxplots). Right panel: Ratios of the lengths of chromosomes between hybrids and pure species. X and Y – sex chromosomes. 2 and 3 – autosomes. Statistical significance was assessed with a two-sample pooled *t*-test.

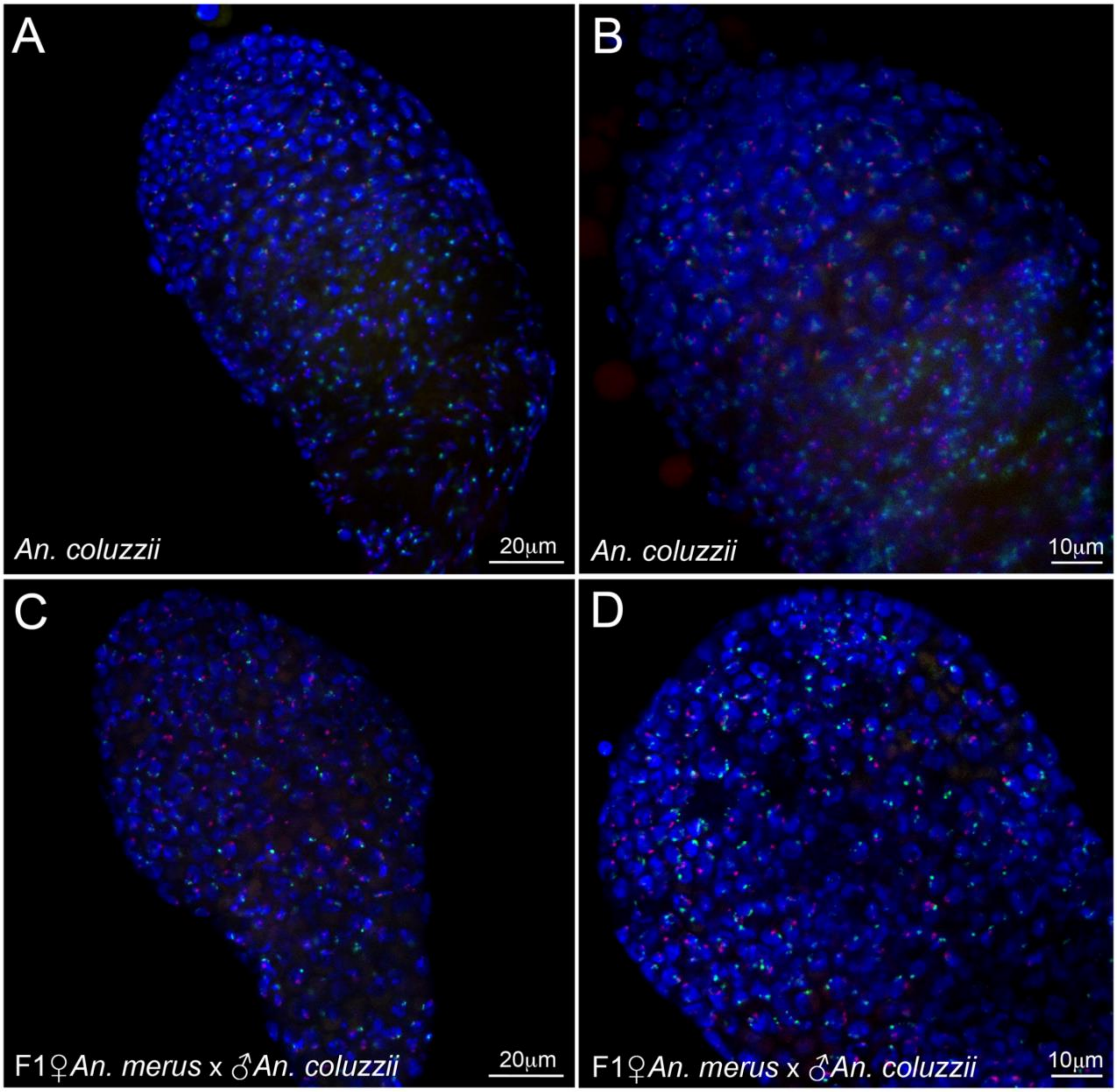


Figure S9. Visualization of pairing between sex chromosomes in testes of a pure species and an interspecies F1 hybrid. (A, B) Testis of a one-day-old adult of *An. coluzzii* MOPTI. (C, D) Testis of a one-day-old adult F1 hybrid of the ♀*An. merus* × ♂*An. coluzzii* MOPTI cross. Whole-mount FISH with 18S rDNA (Cy3), a satellite from Contig_240 (Cy3), satellite AgY53B (Cy5), and satellite AgY477-AgY53B junction region (Cy5) is performed to detect the X (red) and Y (green) chromosomes. Chromatin is counterstained with DAPI (blue).

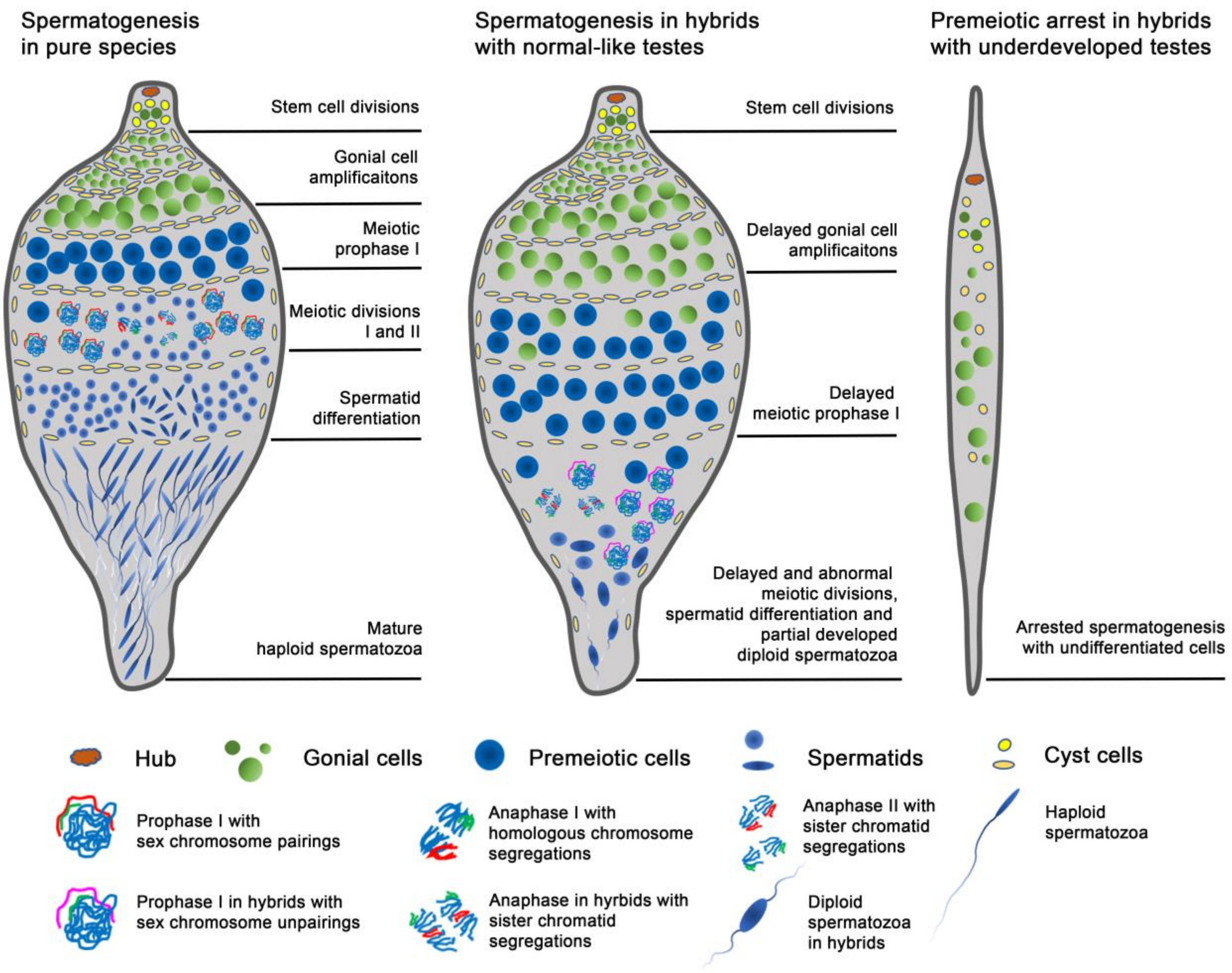


Figure S10. Schematic representation of testis development in pure species and in reciprocal F1 hybrids of the *An. gambiae* complex.
